## Supplementary material for "CBEA: Competitive balances for taxonomic enrichment analysis": S1 File

February 15, 2022

#### 1 Addressing variance inflation due to correlation

In order to perform inference with CBEA, we estimated the null distribution empirically. This can be done either non-parametrically by constructing a null distribution through computing scores on multiple permutations of the data, or parametrically via estimating parameters of a distribution using the same permuted scores. This means that our null distribution for a given set is equivalent to scores computed for sets of similar sizes but containing randomly chosen taxa. We chose two distributional forms for the null: the normal distribution and a two-component mixture normal distribution. For the normal distribution, we estimated parameters using the maximum likelihood using the *fitdistrplus* package [1]. For the mixture normal distribution, we utilized the expectation-maximization procedure from the package *mixtools* [2].

An advantage of estimating the null using a parametric approach is the ability to correct for variance inflation due to inter-taxa correlation within the set compared to overall background correlation [3]. CBEA addresses this issue by combining the location (or mean) estimate from the column permuted raw score matrix with the spread (or variance) estimate from the original un-permuted scores. This still allows us to leverage the null generated via column permutation while using the proper variance estimate taken from scores where the correlation structure has not been disrupted. As such, this procedure assumes that the variance of the test statistic under the alternate hypothesis is the same as that of the null.

For the normal distribution, it is straightforward to combine mean and variance estimates from the respective raw score matrices. For the mixture normal distribution, however, due to the fact that the distribution is made of component-wise means, variances and mixing coefficients, we decided to take an optimization based approach to identifying the component wise variances $\sigma_1$ and $\sigma_2$ such the overall mean and variance estimates come from the respective raw cILR score matrices as detailed above. We can write the optimization problem as follows:

$$\begin{aligned} \min_{\sigma_1,\sigma_2} \quad & \sqrt{\left(\sqrt{(\sigma_1 + \mu_1' - M')\lambda_1' + (\sigma_2 + \mu_2' - M')\lambda_2'} - SD\right)^2} \\ \text{s.t.} \quad & \sigma_1 \geq 10^{-5}, \sigma_2 \geq 10^{-5} \\ \text{where} \quad & M', SD, \mu_1', \mu_2', \lambda_1', \lambda_2' \text{ are constants} \end{aligned} \tag{1}$$

$\lambda_1'$, $\lambda_2'$, $\mu_1'$, $\mu_2'$, and $M'$ are component-wise mixing coefficients, component-wise means, and overall mean of the mixture distribution estimated from column-permuted scores while $SD$ is the overall standard deviation of the mixture distribution estimated from unpermuted scores. We solve for this optimization problem using a quasi Newton method with box constraints (L-BFGS-B) with the default finite-difference approximation of the gradient, as implemented in the *optim* function in R.

There are also different variations to this approach. We can choose to vary both $\sigma_1$ and $\sigma_2$ or to keep either $\sigma_1$ or $\sigma_2$ constant and varying the remaining component. We can assume that null distribution is a two-component mixture distribution where there is one major component with smaller mean (representing the bulk of the distribution centered at the permuted mean) and one minor component with higher mean (representing the inflated right tail). Under this assumption, we can modify the optimization problem to only estimate the variance parameter of the smaller component (i.e. without loss of generalizability keeping $\sigma_1$ constant where $\lambda_1' > \lambda_2'$). This allows for the optimization procedure to more properly capture the right tail distribution rather than increasing weight on the left tail of the distribution which impacts the computation of p-values for a one-sided test. Empirical experiments (data not shown) done on simulated data and random set analyses suggest that this adjustment improves performance of the adjusted CBEA under mixture-normal assumption. In the R implementation of CBEA, users can control this behaviour by specifying the *fix_comp* parameter as part of the *control* argument.

### 2 Numerical Simulations

#### 2.1 Design

Even though real data evaluations provide good estimates for performance of CBEA under typical analysis tasks, it does not allow for understanding of the behavior of the model under different effect sizes, correlations, and sparsity. As such, we also perform parametric numerical simulations by generating microbiome count data under the assumption that it follows a zero-inflated negative binomial distribution, which is a good fit for real microbiome relative abundance data [4]. Suppose $X_{ij}$ are observed counts for a sample $i$ and taxon $j$, then we have the following probability model

$$\mathbf{X}_{ij} = \begin{cases} 0 & \text{with probability } p_j \\ \mathbf{NB}(\mu_j, \phi_j) & \text{with probability } 1 - p_j \end{cases} \tag{2}$$

where $\mu_j$ and $\phi_j$ are mean and dispersion parameters, respectively. To incorporate a flexible correlation structure into our simulation model, we utilized the NorTA (Normal to Anything) method [5]. Given an $n$ by $p$ matrix of values $\mathbf{U}$ sampled from multivariate normal distribution with correlation matrix $\rho$, we can generate target microbiome count vector $\mathbf{X_{.j}}$ for taxa $j$ following the marginal distribution $\mathbf{NB}$ characterized by the negative binomial cumulative distribution function $\mathbb{F}_{\mathbf{NB}}$:

$$\mathbf{X}_{.j} = {\mathbb{F}_{\mathbf{NB}}}^{-1}(\Phi_{U_i}) \tag{3}$$

In this instance, for each taxon $j$, we set elements in $\mathbf{U}_{.j}$ to be zero with probability $p_j$ and applied $\mathbf{NB}^{-1}(\mu_j, \phi_j)$ on non-zero elements to generate our final count matrix $\mathbf{X}$. To ensure that our simulations match closely to real data, we fitted negative binomial distribution using a maximum likelihood approach (with the *fitdistrplus* package in R [1]) to non-zero counts for each taxon from 16S rRNA profiling of stool samples from the Human Microbiome Project (HMP). We take the median values of the estimated mean ($\mu_j$) and dispersion parameters ($\phi_j$) as the baseline of our simulations. For simplicity, we assumed that inter-taxa correlation follows an exchangeable structure with correlation equals to $\rho$.

##### 2.1.1 Simulation scenarios for enrichment analysis at the sample level

To assess type I error rate and power for enrichment significance testing at the sample level, we simulated data based on the schema above, and assessed enrichment for one focal set. Type I error was obtained under the global null as the number of samples where the null hypothesis was rejected at $\alpha = 0.05$ over the total number of samples (which represents the total number of hypotheses tested). Power was obtained using the same formulation as type I error rate but under the global alternate. We treated type I error and power as estimates of binomial proportions and utilized the Agresti-Couli [6] formulation to calculate 95% confidence intervals. Across both analyses, we varied sparsity levels ($p = 0.2, 0.4, 0.6$) and inter-taxa correlation within the set ($\rho = 0, 0.2, 0.5$). For type I error analysis, we also varied the size of the set (50, 100, 150). For power analyses, set size was kept constant at 100 but different effect sizes (fold change of 1.5, 2, and 3). All sample sizes were set at 10,000.

For classifiability, we evaluated the scores against the true labels per sample (indicating the sample has a set with inflated counts) using the area under the receiving operator curve (AUROC). This is a strategy used in Frost [7] which evaluates the informativeness of scores by assessing the relative ranking of samples (i.e.

whether samples with inflated counts are highly ranked using estimated scores). DeLong 95% confidence intervals for AUROC [8] were obtained for each estimate. Simulation settings for classification performance were identical to power analyses as detailed in the previous paragraph.

###### 2.1.2 Simulation scenarios for enrichment analysis at the population level

To assess type I error rate and power for inferece at the population level, we simulated data based on the schema above, and assessed the enrichment of 50 sets (with 100 taxa per set) across 10 replicates per simulation condition. Type I error is calculated as the number of enriched sets over the total number of sets for each simulation under the global null. Power is defined similarly, but instead under the global alternate hypothesis. Estimates and confidence intervals for type I error and power are calculated as cross-replicate mean and standard error. Across both analyses, we varied sparsity levels ($p = 0.2, 0.4, 0.6$), and inter-taxa correlation within the set ($\rho = 0, 0.2, 0.5$). For power analyse, we defined an enriched set as a set where all taxa within a set have inflated means of the same effect size. Half of the sets are defined as enriched across case/control status with varying effect sizes (fold change of 1.5, 2, and 3). Due to the compositional nature of microbiome taxonomic data, simple inflation of raw counts would cause an artificial decrease in the abundance of the remaining un-inflated sets. As such, we applied a compensation procedure as described in Hawinkel et al. [9] to ensure the validity of simulation results. All sample sizes were set at 500.

###### 2.1.3 Simulation scenarios for downstream prediction

To assess predictive performance, we generated predictors based on the simulation schema presented above and evaluated prediction for both binary and continuous outcomes using a standard random forest model [10]. For binary outcomes, we use AUROC similar to the classification analyses above. For continuous outcomes, we used root mean squared error (RMSE). All predictive model fitting was performed using *tidymodels* [11] suite of packages. Across both learning tasks, we varied sparsity ($p = 0.2, 0.4, 0.6$), and inter-taxa correlation ($\rho = 0, 0.2, 0.5$). Continuous outcomes $Y_{cont}$ were generated as linear combinations of taxa counts.

$$Y_{cont} = f(\mathbf{X}) + \epsilon \tag{4}$$

where $\epsilon \sim N(0, \sigma_\epsilon^2)$ and $f(\mathbf{X}) = \beta_0 + \mathbf{X}\beta$. For each simulation, we set $\beta_0$ to be $\frac{6}{\sqrt{10}}$ similar to [12]. The degree of model saturation (the number of non zero $\beta$ values) were varied between 0.1 and 0.5, and signal to noise ratio (SNR $= \frac{\sigma(f(\mathbf{X}))}{\sigma_\epsilon}$) was varied between 1.5, 2, and 3.

For binary outcomes, we generate $Y_{binary}$ as Bernoulli draws with probability $p_{binary}$, where

$$p_{binary} = \frac{1}{1 + \exp(f(\mathbf{X}) + \epsilon)} \tag{5}$$

To ensure a balance of classes, we applied the strategy described in Dong et al. [13] where the associated $\beta$ values are evenly split between positive and negative associations. All data sets generated from prediction tasks have 2,000 samples with 5,000 taxa over 50 sets with a size of 100 taxa per set.

##### 2.2 Results

###### 2.2.1 Statistical Inference

Fig S1 demonstrate type I error evaluations for sample-level inference with CBEA compared to the Wilcoxon rank sum test, which uses a rank-based statistic to compare the mean count difference between taxa in the set its complement. All methods demonstrate good type I error control at $\alpha = 0.05$ under zero correlation across all simulation conditions. However, under both medium ($\rho = 0.2$) and high ($\rho = 0.5$) correlation settings, both the Wilcoxon test and unadjusted CBEA variants show high levels of inflated type I error, where Wilcoxon test performed the worst. On the other hand, adjusted CBEA methods (under both distributions) control for type I error at the appropriate $\alpha$ level even at high correlations. This is opposite from our real data evaluations, where adjusted CBEA demonstrated inflated type I error under random set evalutions.

We also assessed the ability to perform inference at the population level using CBEA similar to GSVA [14]. Here, we test for enrichment of sets across case/control status by generating CBEA scores and performing

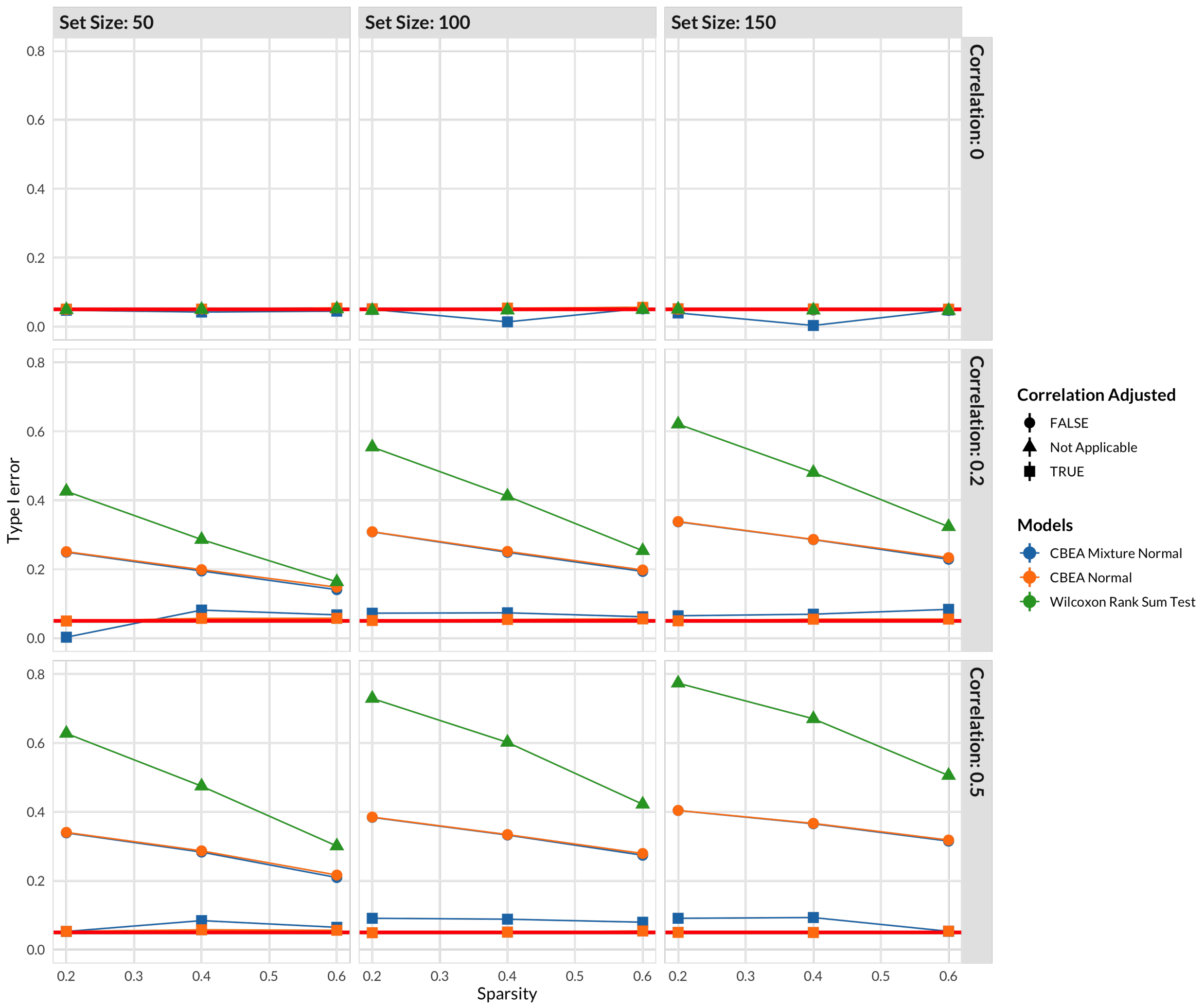


**Figure S1. Simulation results for type I error evaluation for CBEA sample-level inference**. Type I error rate ($y$ axis) was estimated for each approach across data sparsity levels ($x$ axis) across different set sizes (horizontal) and inter-taxa correlation within the set (vertical). We compared variatns of CBEA against a Wilcoxon rank sum test at $\alpha$ of 0.05. For each scenario, a data set of 10,000 samples (equivalent to 10,000 hypotheses) was utilized. Confidence bounds were obtained using Agresti-Couli [6] approach.

Welch's t-test as a difference in means test. We compared the performance of this approach with CBEA and two commonly used methods for differential abundance testing in the microbiome literature: DESeq2 [15] and corncob [16]. Fig. S2 present results for simulation studies for both type I error evaluations. All methods were able to control for type I error across both sparsity and correlation levels, where medium level sparsity ($p = 0.4$) and correlation ($\rho = 0.2$) showed the strongest performance. In these scenarios, the CDF values of CBEA generated under the adjusted mixture normal distribution performed the best. This is different than our real data evaluations, where corncob and DESeq2 showed increased type I error.

##### 2.2.2 Phenotype relevance

We assessed phenotype relevance similar to the main manuscript by assessing statistical power and score rankings via AUROC. Results for this analysis is shown in Fig S3. For statistical power (panel A), under low-correlation settings, all CBEA approaches demonstrate similar power, with the unadjusted methods

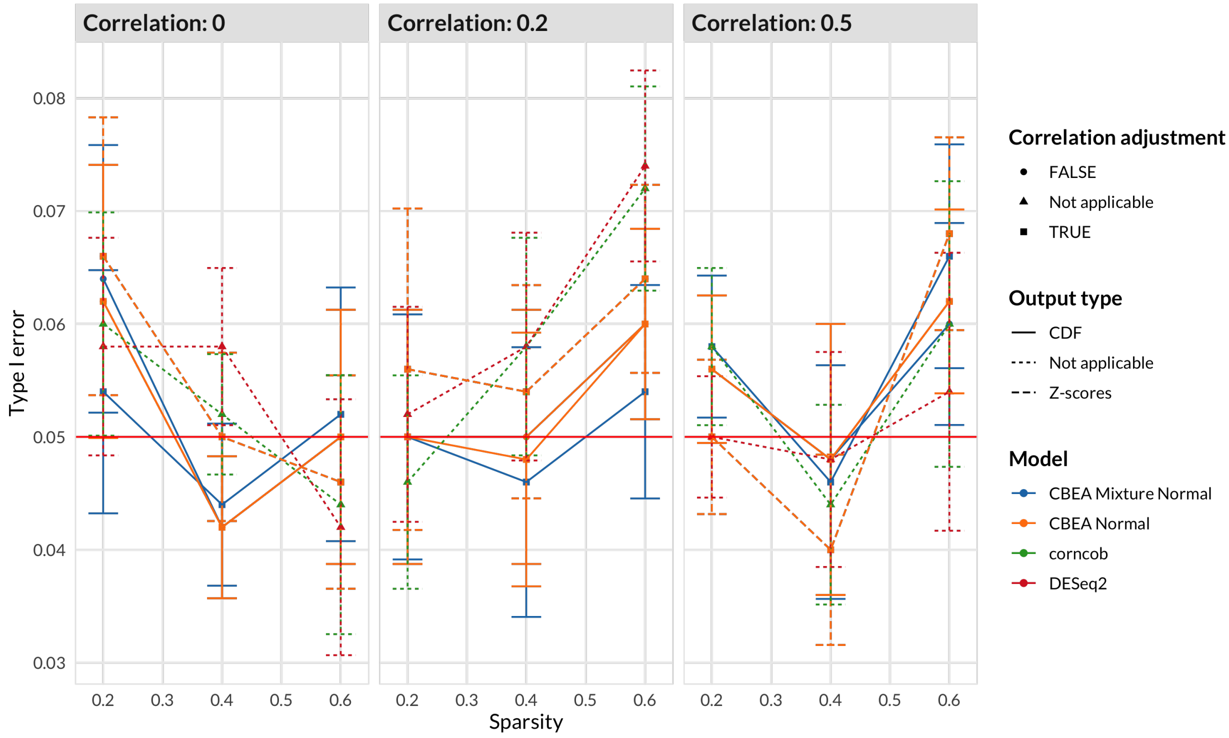


**Figure S2. Simulation results for type I error evaluation for CBEA population-level inference**. Type I error ($y$-axis) was estimated as the average proportion of sets with significant enrichment at 0.05 across 10 replications per simulation condition under the global null. Error bars were estimated using standard errors computed across 10 replicated data sets. Performance was evaluated across different sparsity ($x$-axis) and inter-taxa correlation levels. For CBEA methods, enrichment analysis was performed using a Welch's t-test across case/control status with single sample scores representing set-based features generated by CBEA (across different output types and distributional assumptions). For corncob and DESeq2, set-based features were constructed using element-wise summations.

being slightly more performant at low effect sizes. Notably, all CBEA variants outperformed the Wilcoxon rank sum test. However, as correlation increases, the adjusted CBEA variants showed much lower power, congruent with the perspective that the adjustment process is conservative and trades off power for type I error control. Even at the highest effect size (fold change of 3 in means) adjusted CBEA does not approach 0.8. For score rankings (panel B), all methods are close together in performance, with the Wilcoxon W statistic being the worst performer. These results are similar to that of our real data evaluations. Notably, performance values were not affected by sparsity and increases to near perfect prediction at higher effect sizes.

For population-level analyses, we assessed phenotype relevance as statistical power to detect sets that were simulated to be significantly enriched. Fig S4 showed these results. As expected, power decreases with increasing sparsity, where the effect was attenuated at lower effect sizes. Correlation did impact power, however the difference was not notable. Interestingly, at lower effect sizes both DESeq2 and corncob has comparable power with CBEA, however as effect size increases, the difference in performance values became more stark. This is different than our real data evaluations, where power was more comparable (with slight advantage to corncob and DESeq2).

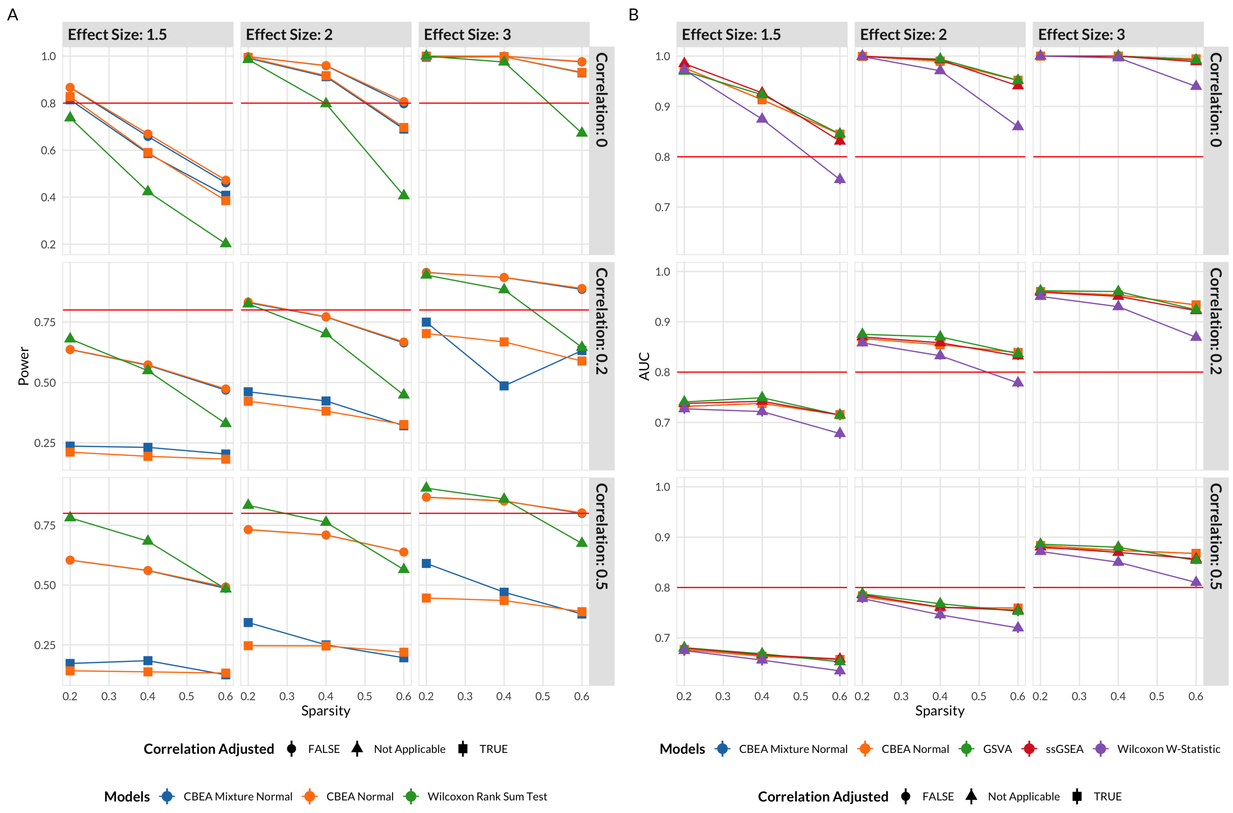


**Figure S3. Simulation results for phenotype relevance valuation for CBEA sample-level inference**. **(A)** demonstrate statistical power ($y$-axis) across different data sparsity levels ($x$-axis) and power **(B)** for differential abundance test across different parametric simulation scenarios. For CBEA methods, differential abundance analysis was performed using a difference in means test (either Wilcoxon rank-sum test or Welch's t-test) across case/control status using single sample scores generated by CBEA (across different output types and distributional assumptions). CBEA associated methods demonstrated similar type I error to conventional differential abundance analysis methods but with more power to detect differences even at small effect sizes.

##### 2.2.3 Prediction analysis

Fig S5 shows results for simulation studies as detailed in the Methods section. Panel A presents results for a regression learning task with a continuous outcome. We observed no difference in performance values across different evaluation models. Overall, prediction error did not change across sparsity levels, and decreases with increasing signal-to-noise ratio (SNR). However, at higher correlation levels, the pattern was more erratic. For example, when $\rho$ is set at 0.5, higher SNR decreases performance only at low sparsity $p = 0.2$, but had the expected pattern medium sparsity $p = 0.4$, where higher SNR correlates with improved performance. This effect seems to be specific to low effect saturation scenarios (only a limited number of taxa sets are associated with the outcome). Interestingly, higher sparsity levels produces better performance.

Panel B represent results for a classification task with a binary outcome with AUROC as the evaluation criteria. Here we observed similar results as that of our real data evaluations, where using CLR transfromed data produces more predictive models across all scenarios. Conversely, GSVA and ssGSEA were consistently under performing when compared to CBEA and CLR. Interestingly, the degree of difference varies across sparsity and inter-taxa correlations. We noticed that increasing sparsity and correlation decreases the gap in performance between CBEA and CLR, while increasing the gap in performance between CBEA/CLR

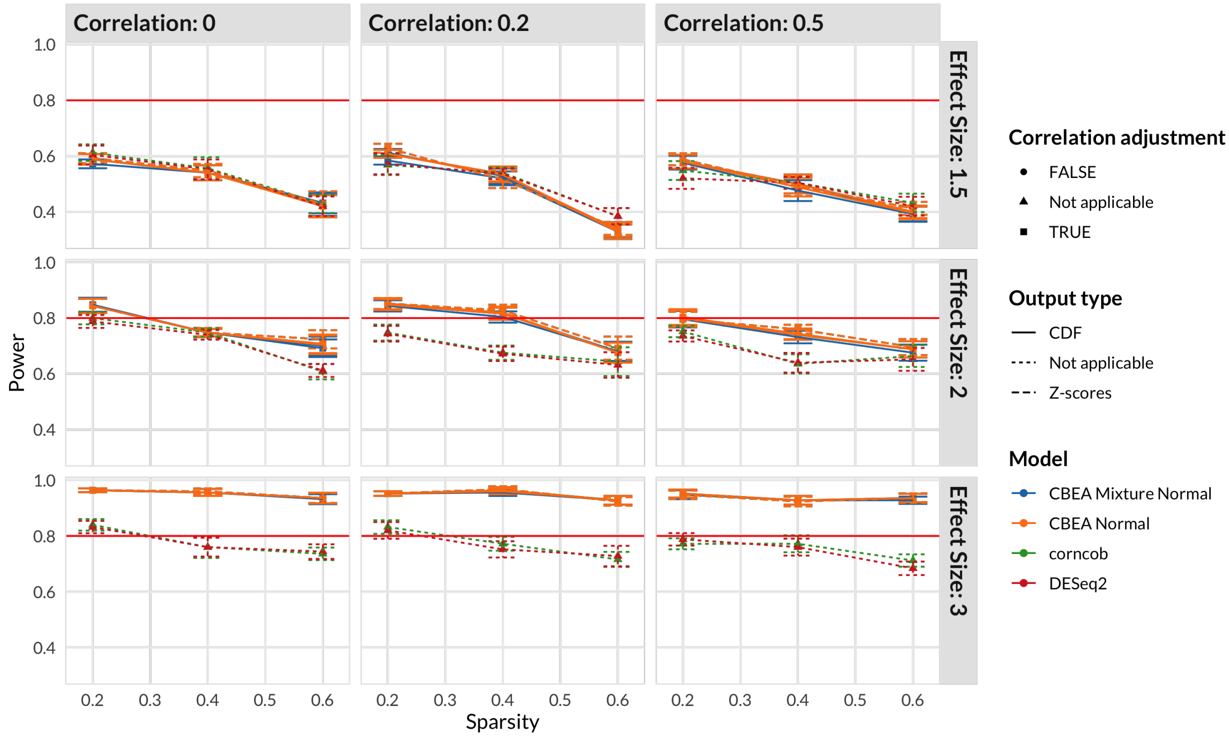


**Figure S4. Simulation results for phenotype relevance evaluation for CBEA population-level inference**. Power ($y$-axis) was estimated as the average proportion of sets correctly identified as significantly enriched (at 0.05) across 10 replications per simulation condition under the global null. Error bars were estimated using standard errors computed across 10 replicated data sets. Performance was evaluated across different sparsity ($x$-axis) and inter-taxa correlation levels. For CBEA methods, enrichment analysis was performed using a Welch's t-test across case/control status with single sample scores representing set-based features generated by CBEA (across different output types and distributional assumptions). For corncob and DESeq2, set-based features were constructed using element-wise summations.

and GSVA/ssGSEA. As such, we can hypothesize that both GSVA and ssGSEA are more sensitive to the degree of inter-taxa correlation and sparsity. Finally, effect saturation did not change model rankings, but did decrease overall performance.

### 3 Runtime

We implemented CBEA in the package *CBEA* on GitHub (`https://www.github.com/qpmnguyen/CBEA`). We evaluated computational time using the *bench* package in R. We applied CBEA to a standard data set generated using our simulation model consisting of 40 sets (of size 20 each) and 500 samples. Benchmark was performed on a single core using a node on the computing cluster (Specifications: Intel Xeon E5-2690 (2.6GHz) with 4GB of RAM).

In general, using the normal distribution is the fastest approach regardless of the total number of permutations performed (2 minutes). However, using the mixture normal distribution increased the runtime many folds, especially for the adjusted approach (highest was 5.53 hours). This time scales with the number of sets evaluated, as well as the number of samples. We believe this is due to the procedure used to estimate parameters of the mixture normal distrubiton as implemented in the *mixtools* package. The default parameters used in CBEA also increased the runtime in order to reduce convergence issues.

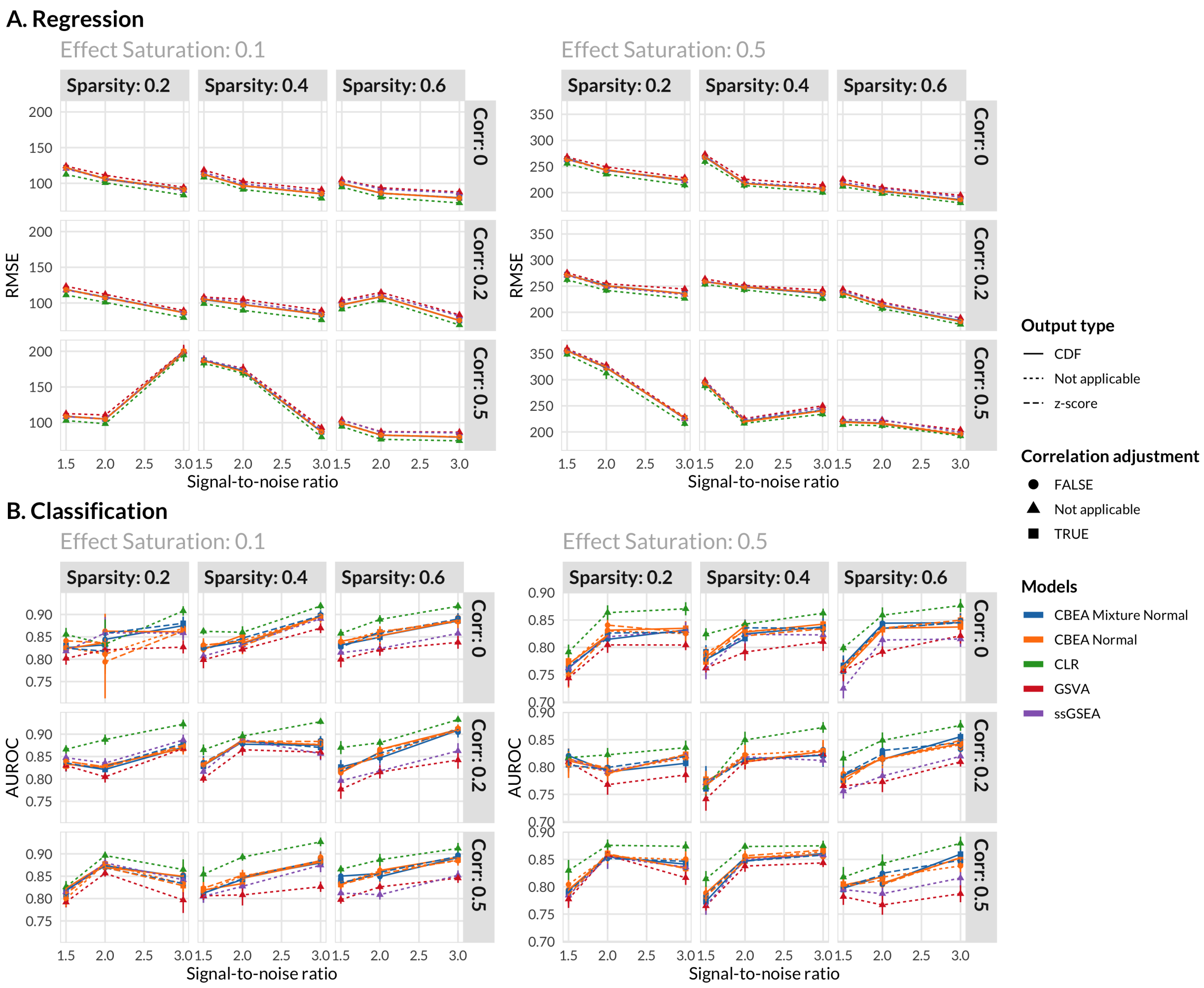


**Figure S5. Simulation results for predictive performance evaluation for CBEA.** Predictive performance of a random forest model (with no hyperparameter tuning) trained on set-based features as inputs. Methods to generate these features include CBEA, ssGSEA, GSVA, and the CLR transformation applied on sum-aggregated sets. Simulation data was generated across different levels of data sparsity, inter-taxa correlation, effect saturation, and signal-to-noise ratio. Panel **(A)** presents performance on a regression task using RMSE (root mean squared error) as the evaluation measure. Panel **(B)** presents performance on a classification task with AUROC as the evaluation measure

In order to reduce runtime, users can attempt the following: Since CBEA fits parametric distributions over permuted values of all samples within a data set (i.e. for $N = 100$ and 10 permutations, the fitting procedure will attempt to estimate parameters from a vector of size 1000 for each set), if the sample size is high users can reduce the number of permutations. Additionally, CBEA also implements a procedure to parallelize computation across sets, which might be applicable to situations where there are a lot of sets to evaluate. Finally, a lot of CBEA approaches work well without parametric fit, so users can use the non-parametric approaches like the permutation test or using raw CBEA scores.

### 4 Single sample type I error diagnostic plot

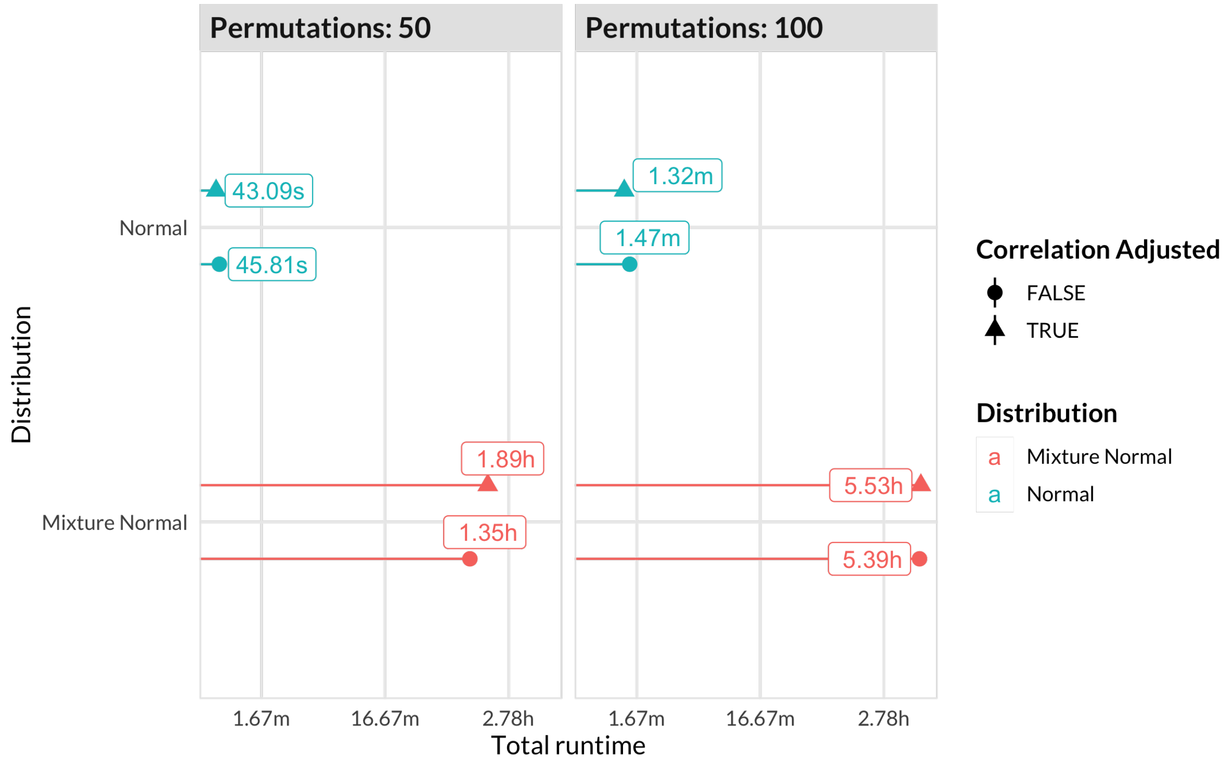


**Figure S6. Runtime performance**. Overall runtime of CBEA under different parameters for a data set of 500 samples, 800 taxa (40 sets of size 20 each). This data set was generated via simulations.

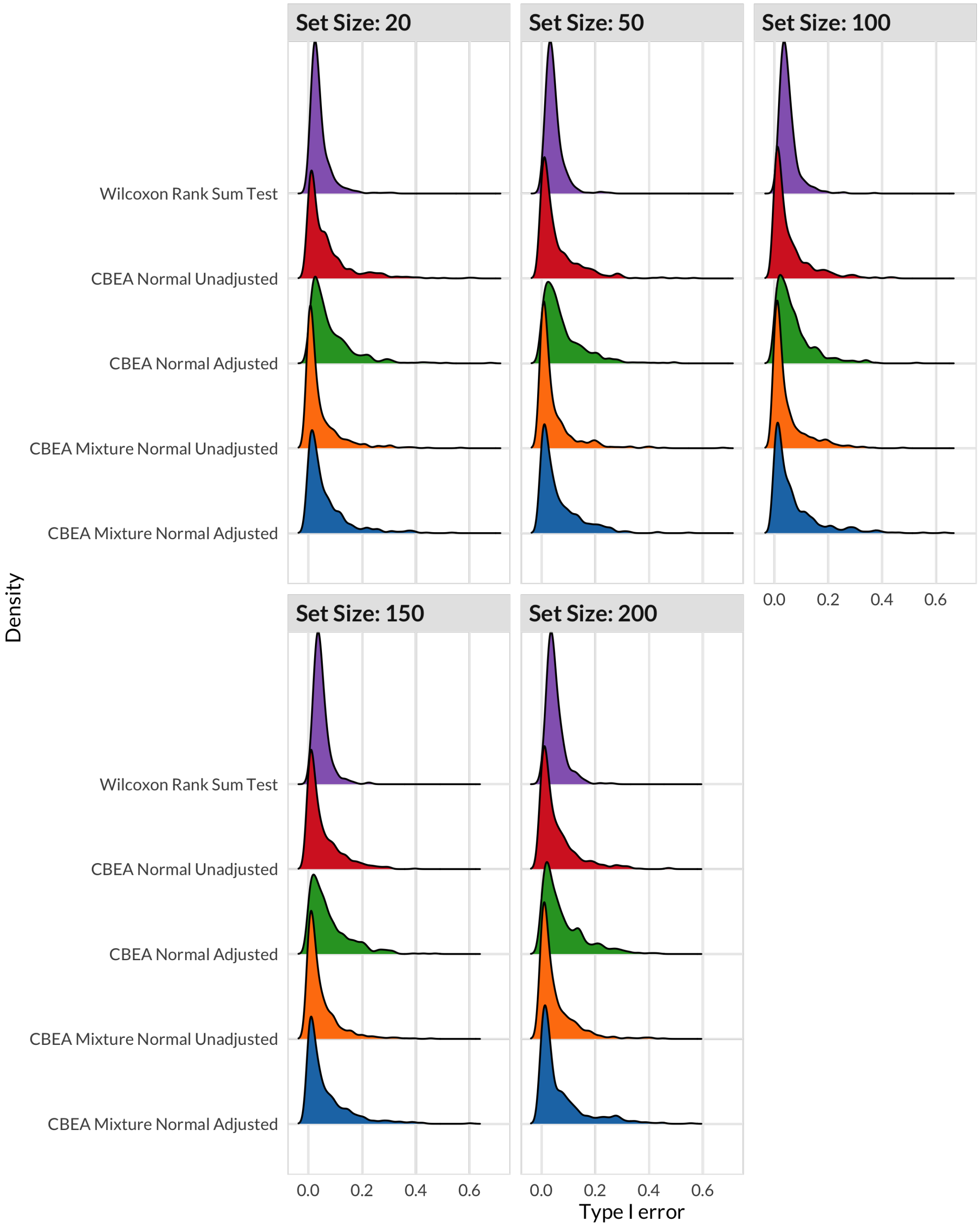


**Figure S7. Distribution of type I error values across app relications in real data random set evaluations for CBEA inference at the sample level**. Density ($y$-axis) for type I error values ($x$-axis) of each evaluated approach for sample-level inference using real data across 500 replications. Here, type I error was estimated as the proportion of samples where a randomly sampled set of different sizes where identified to be statistically significant at $p$-value threshold of 0.05

### References


[1] Delignette-Muller ML, Dutang C. Fitdistrplus: An R Package for Fitting Distributions. Journal of Statistical Software. 2015;64(4):1–34.

[2] Benaglia T, Chauveau D, Hunter DR, Young D. Mixtools: An R Package for Analyzing Finite Mixture Models. Journal of Statistical Software. 2009;32(6):1–29.

[3] Wu D, Smyth GK. Camera: A Competitive Gene Set Test Accounting for Inter-Gene Correlation. Nucleic Acids Research. 2012;40(17):e133. doi:10.1093/nar/gks461.

[4] Calgaro M, Romualdi C, Waldron L, Risso D, Vitulo N. Assessment of Statistical Methods from Single Cell, Bulk RNA-seq, and Metagenomics Applied to Microbiome Data. Genome Biology. 2020;21(1):191. doi:10.1186/s13059-020-02104-1.

[5] Cario MC. Modeling and Generating Random Vectors with Arbitrary Marginal Distributions and Correlation Matrix; 1997.

[6] Agresti A, Coull BA. Approximate Is Better than "Exact" for Interval Estimation of Binomial Proportions. The American Statistician. 1998;52(2):119–126. doi:10.2307/2685469.

[7] Frost HR. Variance-Adjusted Mahalanobis (VAM): A Fast and Accurate Method for Cell-Specific Gene Set Scoring. Nucleic Acids Research. 2020;48(16):e94–e94. doi:10.1093/nar/gkaa582.

[8] DeLong ER, DeLong DM, Clarke-Pearson DL. Comparing the Areas under Two or More Correlated Receiver Operating Characteristic Curves: A Nonparametric Approach. Biometrics. 1988;44(3):837–845.

[9] Hawinkel S, Mattiello F, Bijnens L, Thas O. A Broken Promise: Microbiome Differential Abundance Methods Do Not Control the False Discovery Rate. Briefings in Bioinformatics. 2019;20(1):210–221. doi:10.1093/bib/bbx104.

[10] Breiman L. Random Forests. Machine Learning. 2001;45(1):5–32. doi:10.1023/A:1010933404324.

[11] Kuhn M, Wickham H. Tidymodels: A Collection of Packages for Modeling and Machine Learning Using Tidyverse Principles.; 2020.

[12] Xiao J, Chen L, Yu Y, Zhang X, Chen J. A Phylogeny-Regularized Sparse Regression Model for Predictive Modeling of Microbial Community Data. Frontiers in Microbiology. 2018;9. doi:10.3389/fmicb.2018.03112.

[13] Dong M, Li L, Chen M, Kusalik A, Xu W. Predictive Analysis Methods for Human Microbiome Data with Application to Parkinson's Disease. PLOS ONE. 2020;15(8):e0237779. doi:10.1371/journal.pone.0237779.

[14] Hänzelmann S, Castelo R, Guinney J. GSVA: Gene Set Variation Analysis for Microarray and RNA-Seq Data. BMC Bioinformatics. 2013;14(1):7. doi:10.1186/1471-2105-14-7.

[15] Love MI, Huber W, Anders S. Moderated Estimation of Fold Change and Dispersion for RNA-seq Data with DESeq2. Genome Biology. 2014;15(12):550. doi:10.1186/s13059-014-0550-8.

[16] Martin BD, Witten D, Willis AD. Modeling Microbial Abundances and Dysbiosis with Beta-Binomial Regression. The Annals of Applied Statistics. 2020;14(1):94–115. doi:10.1214/19-AOAS1283.
